# Network-Based Analysis of Human Astrocytes Links Aging to Neurodegenerative and Cardiovascular Diseases

**DOI:** 10.1101/2025.11.05.686746

**Authors:** Patricia M. Bota, Pol Picón-Pagès, Hugo Fanlo-Ucar, Saja Almabhouh, Oriol Bagudanch, Melisa E. Zeylan, Simge Senyuz, Patrick Gohl, Rubén Molina-Fernández, Narcis Fernandez-Fuentes, Eduard Barbu, Raul Vicente, Stanley Nattel, Angel Ois, Albert Puig-Pijoan, Jordi Garcia-Ojalvo, Ozlem Keskin, Attila Gursoy, Francisco J. Muñoz, Baldomero Oliva

## Abstract

Astrocytes are central to brain homeostasis, supporting neuronal metabolism, synaptic activity, and the blood–brain barrier. With aging, these glial cells undergo molecular and functional changes that weaken support functions and promote neuroinflammation, contributing to neurodegeneration. Yet the systems-level mechanisms of astrocytic aging remain poorly defined in human models. Because aging also heightens risk for cardiovascular disease, cognitive impairment, type 2 diabetes, and systemic inflammation, clarifying shared astrocytic pathways is critical for understanding brain–body crosstalk. Using an in vitro human astrocyte model exposed to sublethal oxidative stress (10 µM H₂O₂), we profiled transcriptomic changes and identified differentially expressed genes across antioxidant defences, proteostasis, transcriptional regulation, vesicular trafficking, and inflammatory signalling. We then performed seven network-prioritization analyses on a curated human protein–protein interactome: one seeded with the astrocyte H₂O₂-responsive genes and six with phenotype-associated gene sets (Alzheimer’s disease, cardiovascular disease, cognitive impairment, type 2 diabetes, oxidative stress, and inflammation). Intersecting the top 5% scoring genes from each run yielded a 127-gene core shared across all seven, enriched for proteostasis, DNA repair, mitochondrial regulation, and telomere and nuclear envelope maintenance. Structure-guided analyses highlighted vulnerable interfaces, including lamin A/C–lamin B1, α-actinin–filamins, 14-3-3 dimers, and aminoacyl-tRNA synthetase assemblies, where pathogenic variants are predicted to destabilize or aberrantly stabilize protein interactions. Structure-based interface predictions also highlight potential interactions between APP–VCP/p97 and p53–14-3-3ζ that link proteostasis and stress signalling. Together, these findings define a conserved astrocytic vulnerability network that may couple neurodegeneration with cardiovascular disease and nominate structurally testable targets for biomarkers and interventions.

## Introduction

Astrocytes, the most abundant glial cells in the brain, are essential for maintaining neuronal health and central nervous system (CNS) homeostasis^1^. They maintain extracellular ionic balance and neurotransmitter homeostasis^1,2^, modulate synaptic activity through gliotransmitter release^3,4^,provide metabolic support to neurons^5^, and help regulate cerebral blood flow while reinforcing the blood–brain barrier^6^. Astrocytes also react dynamically to neural injury or disease by entering reactive states and releasing signalling molecules^1^. In neurodegenerative conditions such as Alzheimer’s disease (AD), astrocytes can become excessively reactive and produce reactive oxygen species (ROS) in response to amyloid-β (Aβ)^7^. These ROS exacerbate neuronal oxidative damage, fuelling a self-perpetuating cycle of neuroinflammation that further impairs astrocytes and neurons^8,9^. Chronic oxidative stress in astrocytes disrupts neurotransmitter regulation and weakens BBB integrity, ultimately amplifying neurodegenerative processes^9^. Notably, astrocyte dysfunction is influenced by systemic vascular factors: cardiovascular pathology can trigger or amplify glial stress responses, placing astrocytes at the nexus of cardiovascular and neurodegenerative processes^10,11^. Cardiovascular disease (CVD) and cognitive impairment (CI) are closely interrelated in the aging population, sharing numerous risk factors and underlying molecular mechanisms^12–14^. CI encompasses conditions from mild cognitive deficits to severe dementia, with AD and vascular dementia being the most prevalent forms^15,16^. CVD can accelerate cognitive decline by chronically reducing cerebral perfusion and triggering neuronal dysfunction^17^. Thus, elucidating the molecular links between CVD and CI is vital for developing novel biomarkers, therapeutic interventions, and preventive strategies. Emerging evidence suggests that aging astrocytes contribute to cognitive deterioration via oxidative stress and inflammatory pathways^18,19^. Moreover, systemic metabolic conditions such as coronary heart disease, atherosclerosis, and type 2 diabetes (T2D), which are strongly correlated with both CI and AD, can directly or indirectly impair astrocytic function^20,21^. This multifactorial convergence implies that astrocytes lie at the intersection of neurodegenerative and systemic aging processes, warranting a systems-level approach to decipher common mechanisms. Protein–protein interaction (PPI) networks provide a powerful framework to integrate diverse molecular data and identify hub proteins that link converging pathological pathways. In this study, we combined transcriptomic profiling with network-based analysis and structural modelling to investigate common mechanisms underlying astrocyte aging. Using a human cortical astrocyte cell-culture exposed to chronic sublethal oxidative stress (10 µM H₂O₂) as an in vitro aging paradigm, we first characterized global transcriptomic changes. We then ran seven separate network-prioritization analyses: one seeded with the astrocyte H₂O₂ response and six seeded with the gene sets for AD, CVD, CI, T2D, I, and OS. In each run, genes were ranked by topological proximity to the seeds, and the top 5% defined the phenotype subnetwork. Finally, we evaluated how disease-associated mutations might disrupt critical protein-protein interfaces. This integrative strategy, spanning from gene expression profiles to systems-level networks to atomic-level interactions, provides a comprehensive view of how astrocytic stress responses may bridge neurodegenerative and cardiovascular disease processes. Our findings reveal conserved molecular vulnerabilities in aging astrocytes and offer mechanistic insights that could inform biomarkers and therapeutic targets for age-related neurological and cardiovascular disorders.

## Materials & Methods

### Cell line

Human cortical astrocytes (HA) were maintained in astrocyte medium (AM; ScienCell) supplemented with 2% fetal bovine serum (FBS), 1% astrocyte growth supplement, and 1% penicillin–streptomycin. Cultures were maintained at 37°C in a humidified incubator with 5% CO_₂_.

### Cell viability assays

Cells were seeded into 96-well plates at a density of 2.5×10^4 cells per well. They were then treated with increasing concentrations of H_2_O_2_ for 24 h. After treatment, 10% of total well volume of 3-(4,5-dimethylthiazol-2-yl)-2,5-diphenyltetrazolium bromide (MTT) stock solution (5 mg/mL in PBS; Sigma) was added to each well and incubated for 2 h. MTT reduction was determined in a plate reader spectrophotometer at 540 and 650 nm. Control cell values were assumed as a 100% of survival.

### Transcriptomic analysis

Cells were seeded in 6-well plates at a density of 5×10^5^ cells/well and left to grow for two days. After washing twice with PBS and once with FBS-free AM medium, cells were treated with 10 μM H_2_O_2_ in 1 mL of FBS-free AM medium for 24 h.

Total RNA was extracted, and its quality and concentration were assessed using an Agilent TapeStation 4200 with RNA ScreenTape. Polyadenylated RNA was captured from 100 ng of total RNA using the NEBNext Poly(A) mRNA Magnetic Isolation Module (New England BioLabs, Ipswich, MA). Stranded RNA-seq libraries were prepared using the NEBNext Ultra II Directional RNA Library Prep Kit for Illumina (New England BioLabs) according to the manufacturer’s instructions. The libraries were amplified by PCR using NEBNext Multiplex Oligos for Illumina (Dual Index Primers Set 1; New England BioLabs). Library quality and size distribution were verified on an Agilent TapeStation 4200 with High Sensitivity D1000 ScreenTape. Finally, libraries were pooled and sequenced (2×75 bp paired-end) on an Illumina NextSeq 500 High Output flow cell.

### Protein–protein interaction network assembly

We built the network of protein–protein interactions (PPI) using BIANA^22^, a Python framework for integrating biological data. First, we retrieved the most recent interaction datasets from IntAct^23^, BioGRID^24^, HIPPIE^25^ and InnateDB^26^, and supplemented them with annotations from UniProt^27^, Reactome^28^, NCBI Gene^29^, Taxonomy, PSI-MI-OBO, GO^30,31^, HGNC^32^, ConsensusPathDB^33^, I2D^34^ and DisGeNET^35^. Following BIANA’s guidelines, we parsed each dataset without altering its original identifiers or metadata. After loading all sources into BIANA’s knowledge database, we unified equivalent entries across resources: proteins sharing a common Entrez Gene ID or, when that was unavailable, both the same Taxonomy ID and identical amino-acid sequence, were merged into single entities. Each consolidated record was then assigned a unique BIANA identifier, effectively reducing redundancy while preserving full traceability. The human-specific protein interaction network (PIN) was retrieved from the unified BIANA database. To reduce biases typically associated with detection methods favouring well studied proteins, we applied stringent filtering criteria, selecting only high confidence interactions validated by relatively unbiased experimental techniques. Specifically, accepted interaction methods included cross-linking studies, classical two-hybrid, two-hybrid arrays, two-hybrid prey pooling validated two-hybrid, two-hybrid pooling, biochemical assays, proximity labelling technology, and enzymatic studies. After filtering, the resulting PPI network comprised 16 009 human proteins connected by 690 067 interactions, which we then used as input for gene prioritization analyses with the GUILD^36^ framework.

### Selection of Genes Associated with Pathophenotypes

Genes linked to specific phenotypes were selected from databases using keyword searches and association data. DisGeNET and OMIM^37^ were employed to retrieve genes associated with AD, T2D, and CI, while gene descriptions in UniProt were used to identify those related to OS and inflammation. For CVD, we used keyword combinations across DisGeNET and UniProt to gather relevant gene associations. This approach ensured comprehensive coverage of genes linked to the targeted aging-associated conditions.

### Prioritization of genes/proteins

We prioritized genes by propagating relevance scores from known disease-associated “seed” genes across the PPI network and integrating the outputs of multiple complementary algorithms into a single consensus ranking. For this purpose, we used GUILD^36^, a genome-wide network-based disease-gene prioritization framework that identifies candidate genes based on their topological proximity to seeds in the interactome. GUILD implements three topology-based algorithms: NetShort, which ranks genes by their weighted shortest-path distances to seed nodes, emphasizing proteins that lie on minimal connection routes; NetZcore, which propagates relevance through immediate neighbours and applies z-score normalization against randomized networks to reduce hub bias; and NetScore, which iteratively diffuses scores along shortest paths from seeds, updating rankings over multiple rounds until convergence. In addition to these local topology-based algorithms, GUILD includes four global diffusion and propagation methods that disseminate relevance across the entire network: PageRank with priors^38^, Functional Flow^39^, Random Walk with Restart^40^, and Network Propagation^41^. We also incorporated DIAMOnD^42^, which extends disease-related modules by iteratively adding proteins based on statistically significant connectivity to existing seeds, calculated via hypergeometric testing. Individual scores from all seven methods were normalized and aggregated into a unified consensus score. In this way, the eight methods capture different aspects of network connectivity – from local shortest-path relationships to global diffusion patterns – and together provide a robust prioritization of candidate disease genes.

### Functional enrichment

We assessed the overrepresentation of biological functions among both the seed proteins and the top-ranked candidate genes. Functional annotations were derived from Gene Ontology (GO) biological process terms and *Reactome* pathways. To ensure high-confidence results, only GO annotations supported by curated or experimental evidence codes (EXP, IDA, IMP, IGI, IEP, ISS, ISA, ISM, ISO) were included. Enrichment was evaluated using a one-sided Fisher’s exact test, comparing the observed overlap between the gene sets and annotated terms against random expectation. To account for multiple testing, p-values were corrected using both the Benjamini-Hochberg^43^ false discovery rate procedure and the Bonferroni correction. Annotations with an FDR-adjusted p-value below 0.05 were considered significantly enriched.

### Structure modelling of PPIs

Experimentally solved binary complexes currently cover less than 5% of known human protein–protein interactions^44^, making it necessary to rely on predicted structures for large-scale interactome analyses. To obtain three-dimensional models for the prioritized protein– protein interactions we used two sources: (1) the Comparative Modelling and Data-Driven Docking (CM2D3)^45^ database and (2) the Protein Interactions by Structural Matching (PRISM)^46^ web server. CM2D3 provides predicted complex models derived by both homology modelling and data-driven docking, whereas PRISM is a knowledge-based PPI prediction server that matches input protein structures to experimentally resolved interface templates. The structures retrieved from CM2D3 were first subjected to energy minimization using FoldX^47^ (v2025.12.31) to correct rotamer and backbone distortions (using the *RepairPDB* command). After repair, the FoldX *AnalyseComplex* routine was applied to each wild-type model to calculate its baseline binding free energy and to delineate interface residues (defined as any residue whose Cα atom lies within 12 Å of the partner chain). Pathogenic missense variants were obtained from ClinVar^48^ (accessed June 2025) and mapped onto UniProtKB canonical sequences to ensure consistent residue numbering. Variants located at the predicted interface (i.e. coinciding with the annotated interface residues) were selected for in silico mutagenesis. Using FoldX’s *BuildModel*, each single-point mutation was introduced into the repaired complex structure, followed by a second AnalyseComplex run to obtain the mutant binding energy. The change in interface stability (ΔΔG_interface) was then computed as the difference between mutant and wild-type binding free energies. Mutations yielding ΔΔG_interface > 1 kcal/mol were classified as destabilizing, in accordance with published FoldX benchmarks. In parallel we used PRISM to search for pairs of target structures whose surface patches resemble the two complementary sides of experimentally resolved template interfaces; candidate complexes are then energy minimized and ranked by a global binding-energy score. PRISM predictions were used to identify structurally plausible interfaces for high-priority interactions that lacked CM2D3 models.

## Results

### 1. Transcriptional response in astrocytes under oxidative stress

To determine the cytotoxic threshold of the oxidative agent in human astrocytes, we assessed cell viability across increasing concentrations (Figure 1). We identified 10 µM as a high but subtoxic concentration to perform the transcriptomics assays. Exposure of human astrocytes to mild oxidative stress induced by subtoxic concentrations of hydrogen peroxide (10 µM H_2_O_2_) leads to a broad transcriptional response involving both protective and potentially damaging pathways. This dual response illustrates the inherent plasticity of astrocytes under stress, with gene expression patterns aimed at restoring cellular balance while also hinting at possible long-term dysfunction if the stress remains unresolved.

**Figure 1.**
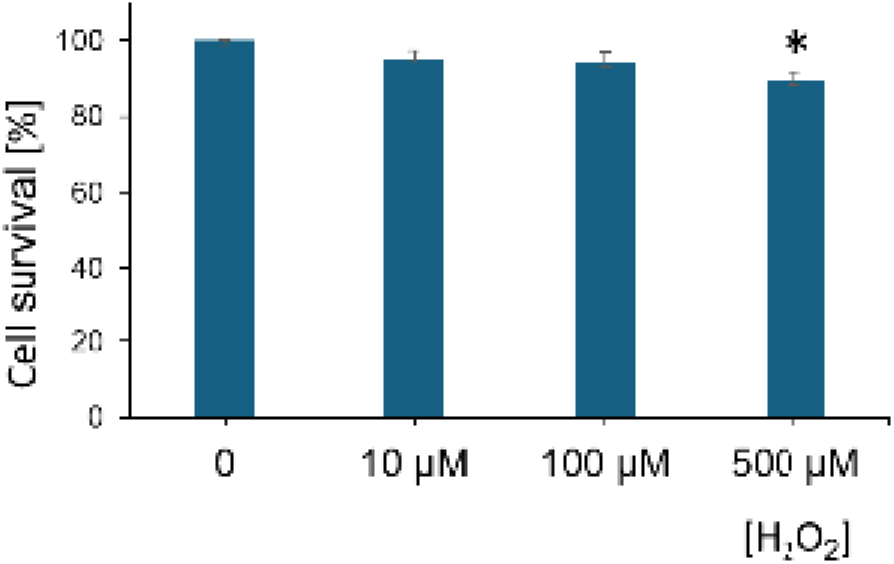
Dose-dependent effect on the viability of human astrocytes in culture. The human astrocyte cell line (HA) was treated with increasing concentrations of H_2_O_2_ for 24h. Cell survival was measured by MTT reduction and expressed as a percentage relative to the control assumed as 100%. Data represent the mean ± SEM of 3 independent experiments performed in triplicate. * p < 0.05 vs. the control, as determined by ANOVA followed by Bonferroni post-hoc test.

A substantial number of upregulated genes reflect the activation of antioxidant defences and maintenance of mitochondrial function (Supplementary Table 1). PRDX5, encoding a mitochondrial peroxiredoxin, detoxifies hydrogen peroxide and organic peroxides and is known to protect cells from oxidative damage^49^. ROMO1 (Reactive Oxygen Species Modulator 1) was also elevated; this inner mitochondrial membrane protein regulates redox signalling and contributes to mitochondrial homeostasis, particularly under oxidative challenge^50^. Increased expression of MIF (Macrophage Migration Inhibitory Factor), a cytokine with both immunomodulatory and oxidoreductase activity, suggests a dual role in coordinating inflammatory responses and enhancing resistance to oxidative injury^51^. Protein quality control mechanisms are also enhanced. STUB1 (E3 ubiquitin-protein ligase CHIP) targets misfolded or damaged proteins for degradation via the ubiquitin–proteasome system and has a known protective role against oxidative stress-induced cell death^52^. Genes such as SERF2, NUDC, and TRMT112 were upregulated, consistent with their roles in stabilizing RNA-protein complexes and supporting proper protein folding, sustaining protein functionality during oxidative stress^53–55^. Simultaneously, we observe increased expression of genes associated with ribosomal function and protein synthesis, such as RPS9, RPL41, POLR3K, and RRP7A. The presence of CTU1, which modifies tRNAs to ensure translational accuracy^56^, further underscores the cell’s attempt to recalibrate its proteome. The data also reveal upregulation of several genes related to inflammatory and signalling pathways. CYBA, a subunit of NADPH oxidase, could amplify oxidative signals and contribute to inflammatory activation^57^. GRK6 and DVL1, involved in GPCR^58^ and Wnt signalling^59^ respectively, play key roles in modulating inflammatory and plasticity-related pathways in the CNS. Increased expression of RAB13 and YIF1B suggests heightened intracellular vesicle trafficking^60,61^, potentially affecting the secretion of cytokines or gliotransmitters. Interestingly, the transcriptional profile includes upregulation of genes related to the cell cycle (e.g., PKMYT1 and MXD4), which may signal stress-induced cell cycle re-entry or glial senescence. The upregulation of EGFL7, an established endothelial-secreted angiogenic factor^62^, indicates astrocytes may modulate vascular homeostasis under oxidative stress by interacting with endothelial extracellular matrix proteins, with potential consequences for blood–brain barrier stability.

Exposure to subtoxic levels of H₂O₂ not only triggers protective mechanisms in astrocytes but also leads to the repression of key genes (Supplementary Table 1), suggesting a reduction in cellular activities critical for homeostasis, signalling, and structural maintenance. The widespread down regulation reflects a coordinated suppression of energy-demanding or complex regulatory pathways, possibly as an adaptive mechanism to conserve resources under oxidative stress, but also potentially contributing to functional decline if the stress persists. A group of repressed genes are involved in transcriptional regulation and chromatin remodelling, such as KLF12, CHD9, ASXL2, JMJD1C, KDM3A, and AFF4. These genes play essential roles in controlling gene expression, epigenetic modifications, and developmental programs. Their downregulation may dampen the astrocytic transcriptional flexibility, limiting the cell ability to respond dynamically to environmental cues. TEAD1, a key transcription factor in the Hippo pathway involved in cell growth and survival^63^, is also repressed, suggesting impaired proliferative or reparative capacity. Genes controlling intracellular transport and cytoskeletal organization are also significantly downregulated. These include VPS13A, VPS13C, ARFGEF1, DYNC2H1, and RANBP2, which govern lipid and vesicle transport as well as nuclear-cytoplasmic trafficking^64–67^. Similarly, DOCK9 and PSD3, involved in cytoskeletal remodelling, may affect astrocyte shape and mobility. A subset of downregulated genes are central to signal transduction and receptor function. For instance, IL6ST (gp130), a co-receptor for IL-6 family cytokines, and LIFR (Leukemia Inhibitory Factor Receptor) are both involved in neuroprotective signalling and gliogenesis^68^. Their repression suggests an impaired ability to respond to neurotrophic or inflammatory stimuli. SOS1, TAOK1, and BMPR2 are kinases or signalling adaptors that regulate pathways like MAPK and BMP, further indicating a broad attenuation of astrocytic signalling^69,70^. A critical concern is the downregulation of DNA repair genes such as ATM, SETX, SMG1, CDK12, and TLK1. This repression could impair genomic integrity and contribute to glial aging^71,72^. The repression of structural and adhesion-related genes like ITGAV, VCAN, DST, HMCN1, and CAMSAP2 may compromise the ability of astrocytes to maintain contact with other cells and the extracellular matrix, impacting BBB maintenance and tissue architecture^73–76^. SLC7A11 (a subunit of the cystine/glutamate antiporter) is downregulated, which may impair glutathione synthesis and reduce antioxidant capacity, counteracting protective responses and promoting oxidative damage. Finally, the downregulation of genes involved in protein degradation and turnover, such as LTN1, UBXN7, and USP34, suggests a reduction in proteostasis maintenance under mild oxidative stress^77–80^. This may lead to accumulation of damaged or misfolded proteins if the stress persists.

Overall, this transcriptional response represents a dual-edged cellular adaptation. On one hand, the upregulation of antioxidant enzymes, chaperones, and other stress-response genes should enhance survival, damage repair, and recovery of function. On the other, the concurrent repression of genes controlling transcription and chromatin regulation, receptor-and kinase-mediated signalling, intracellular transport and cytoskeletal/adhesion structures, DNA repair, glutathione and redox metabolism, and protein degradation indicates a strategic downscaling of activity. This likely helps prioritize defence in the short term but also creates a vulnerability: if oxidative stress becomes chronic, impaired neuroprotective, redox, and structural support functions in astrocytes may promote pro-inflammatory and degenerative changes, contributing to disease progression in conditions such as Alzheimer’s disease and vascular cognitive impairment.

### 2. Analysis of networks associated with selected phenotypes

To place the astrocyte transcriptomic changes in disease context, we mapped the H₂O₂-responsive genes onto a human protein–protein interaction (PPI) network. Each differentially expressed gene served as a seed, and we propagated influence across the interactome with the GUILD framework to identify proteins tightly connected to the astrocyte response. We then repeated this propagation for specific disease phenotypes (AD, CVD, T2D, CI, inflammation, OS) using their associated genes as additional seeds. Consensus across multiple topology-based algorithms yielded, for each phenotype, a prioritized subnetwork enriched for nodes central to disease biology and potentially influenced by astrocyte signals. This systems-level analysis reveals shared molecular players and pathways through which astrocyte oxidative stress may couple to neurodegenerative and cardiometabolic pathology.

### 2a. Network of Proteins Affected by Oxidative Stress in Astrocytes (PAOSA)

Using the top 5% of nodes prioritized by GUILD from astrocyte over- and under-expressed seeds, we observed coherent but opposing cellular programs. The overexpressed–seed subnetwork was enriched for protein homeostasis and translation (GO: cytoplasmic translation, translational initiation), proteostasis safeguards (chaperone-mediated protein folding, protein stabilization), ubiquitin-dependent quality control (regulation of protein ubiquitination), and RNA processing (mRNA splicing), with additional signals indicating feedback restraint of stress pathways (negative regulation of stress-activated MAPK cascade) and a telomere-maintenance module. In contrast, the downregulated–seed subnetwork showed broad repression of gene-regulatory capacity, with strong enrichments for RNAPII-mediated transcriptional regulation (both positive and negative), chromatin remodelling and nucleosome disassembly, and regulation of miRNA transcription. The full list of prioritized genes and enriched pathways is provided in GitHub (see Data & Code Availability). Together, these network-level enrichments substantiate a dual response to mild H₂O₂ in astrocytes. We observe activation of proteostatic and translational control programs alongside a coordinated inhibition of transcriptional/epigenetic machinery, supporting the protective-versus-vulnerable balance described above.

### 2b. Network of Proteins of patho-phenotypes related to aging

The AD subnetwork recovers the canonical seeds (APP, MAPT, PSEN1/2, APOE, BACE1, CLU) and established risk/pathway genes (PICALM, BIN1, SORL1, CR1/CD33, IDE, IGF1R/INSR, BDNF, GSK3B), confirming disease specificity. The functional enrichment analyses supports amyloid-β biology^81^ (cellular response to Aβ; negative regulation of Aβ formation; amyloid fibril/fibre formation), ER proteostasis^82^ (positive regulation of ERAD; calnexin/calreticulin cycle; HSF1 activation) and mitochondrial protein quality control (UPRmt; mitochondrial protein degradation)^83^, nitric-oxide biosynthetic regulation^84^, and insulin/IGF-driven signalling converging on PI3K–AKT and MAPK/ERK^85^. Functional enrichment using *Reactome* paths further reinforces IGF1R–SHC and insulin-receptor cascades; activated NTRK2/TrkB via FRS2/3 with PI3K, PLCG1, and RAS branches; FOXO-mediated transcription; and insertion of tail-anchored proteins into the ER, integrating metabolic, synaptic-signalling (including NMDA receptor–RAS), vascular-NO, and proteostasis dimensions of AD. Additional supported signals include endothelial programs (VEGFR2/Tie2 with PECAM1 interactions), AGE–RAGE signalling, and CDK5-linked neurodegenerative pathways. Selenoamino-acid metabolism is also enriched.

The T2D subnetwork is driven by hallmark metabolic regulators: INSR, IRS1, AKT2, PPARG, TCF7L2, GCK, SLC30A8, HNF4A, IGF2BP2, PPARGC1A, SREBF1, LEPR, together with the seeded nodes LMNA (lamin A/C; nuclear-envelope and chromatin organization shaping metabolic gene programs) and TP53 (stress-responsive checkpoint integrating redox, mitochondrial, and apoptotic control). GO terms highlight glucose homeostasis, cellular response to insulin, insulin receptor signalling, positive regulation of D-glucose import, regulation of insulin secretion, leptin-mediated signalling, oxidative-stress responses, and regulation of extrinsic apoptotic signalling via death-receptor pathways. *Reactome* paths reinforce this axis with insulin-receptor and IRS activation, SHC/IGF1R events^86^, FOXO-mediated transcription^87^, transcriptional regulation of white adipocyte differentiation^88^, leptin signalling, and endothelial mechanotransduction through PIEZO1/integrins.

The CVD subnetwork brings together circadian clock genes (CLOCK, PER1/2, CRY1/2, NPAS2) with vascular and metabolic regulators (NOS3, ACE, AGT, VCAM1, ICAM1, VWF, APOB/APOE, LPL, ADRB1) and signalling nodes (AKT1, PTEN, PIK3CA, BDNF, SORT1), supporting endothelial function, lipoprotein handling, and circadian control of cardiovascular physiology. Enrichment signals converge on circadian control tightly linked to endothelial biology: clock programs^89^ couple to disturbed-flow sensing through PIEZO1–integrin pathways and to nitric-oxide production^90^, shaping vascular tone. Lipoprotein–platelet crosstalk is prominent, with LDL-driven platelet sensitization and chylomicron assembly/remodelling pointing to altered lipoprotein handling in the vessel wall^91^. We also see a dampening of glucocorticoid-receptor signalling and engagement of the angiopoietin– Tie2 axis, consistent with shifts in endothelial quiescence versus activation^92^. In parallel, NF-κB/IL-1 inflammatory activity^93^ and strong cell-cycle/DNA-damage/senescence signatures indicate endothelial stress and turnover.

The inflammation subnetwork is anchored by the seed FLNC (filamin-C; an actin crosslinker that transduces mechanical forces at adhesion sites and organizes cytoskeletal signalling), alongside canonical innate–adaptive mediators IL1B, TNF, IL10, TLR2/4 (and TLR3), NFKB1/RELA/NFKB2, MYD88, CASP1/PYCARD, TBK1, PPARG, VCAM1/ICAM1, MMP2/9, PARP1, HMGB1, and HSPA1A/B, defining a strong inflammatory and stress-response core. Enrichment highlights canonical and non-canonical NF-κB activation; up-regulation of IL-1β/IL-6/IL-8^94^; TLR signalling including TLR2:TLR6 and LPS responses; PI3K–AKT and ERK/JNK cascades; chemokine production; endothelial migration and vascular remodelling; antibacterial defence; regulation of superoxide and nitric oxide^95^; LUBAC-mediated linear polyubiquitination^96^; and modulation of extrinsic apoptotic signalling.

The CI subnetwork is dominated by peroxisome-biogenesis genes (multiple PEX family members) and neuronal/repair factors (APP, ATXN1, FLNA, UBA1), together with mitochondrial and DNA-repair modules (SDH subunits, ERCC2/3/4/5, XPA/XPC, ATR, APTX, XRCC4) and signalling nodes (INSR, AKT1, PIK3CA, PTEN). Enrichments focus on peroxisome organization and matrix import (receptor recycling, substrate release, docking, translocation, membrane import), nucleotide-excision repair and homologous recombination, cellular responses to reactive oxygen species, and pexophagy, supporting a CI network centred on peroxisome biogenesis/turnover^97^ and DNA-damage repair^98^.

The OS subnetwork recovers a core redox/stress architecture: NFE2L2/NRF2, SOD1/2, CAT, PRDX1–6, TXNRD1, GSR, PARK7, G6PD, HMOX2, together with mitochondrial quality-control, DNA-repair, proteostasis, and survival regulators (TP53, MAPK14, AKT1/2, FOXO1, HSPA1A/B, HSPB1, PARP1, APEX1, RAD51/XRCC3/ERCC3, PRKN/PINK1, PRDX3/5). Enrichment profiles emphasize cellular and organismal responses to oxidative stress and hydrogen peroxide, suppression of ROS-induced intrinsic (neuronal) apoptosis, base-excision and transcription-coupled nucleotide-excision repair, DNA-damage responses^99^, regulation of mitochondrial membrane potential and organization, induction of autophagy^100^, negative regulation of ROS metabolism, protein repair, H₂O₂ and superoxide detoxification, negative TOR signalling, ubiquitination control, and PERK-mediated UPR.

Interestingly, a small set of non-seed genes consistently achieve high ranks. Most notably, TGOLN2 (TGN46) and the RPA1–3 complex, and in several networks LMNA, recur as prominent hubs. This behaviour aligns with network-propagation theory: algorithms such as NetShort, NetScore, NetZcore, and diffusion methods prioritize nodes that lie on many short paths between seed genes and that connect multiple functional modules. Functionally, TGOLN2 occupies the trans-Golgi sorting/trafficking nexus influencing secretory and membrane-protein routing; RPA1-3 anchors DNA replication/repair and replication-stress responses; and LMNA contributes to nuclear-envelope and chromatin integrity. Their high connectivity and central placement in the interactome therefore make them robust cross-phenotype bridges, explaining why they rank highly even without being seeded a priori.

### 2c. Overlap between the network of PAOSA and the networks of selected phenotypes

Intersecting the top 5% GUILD-prioritized genes from the overexpressed PAOSA with those from AD, CVD, T2D, OS, inflammation, and CI uncovered a large shared-core. As shown in Figure 2, 127 genes are shared by the seven networks, an unexpectedly large intersection that points to a common molecular backbone across aging-related conditions. Consistency across methods was high as most shared genes were recovered by multiple GUILD algorithms (Figure 3), underscoring prioritization robustness. External validation against GenAge^101^ identified 28/127 aging-annotated genes (hypergeometric p-value ≈ 4.8×10⁻²²), supporting the biological relevance of this core. The complete ranked gene lists for each phenotype, including consensus scores, are available in the GitHub repository (see Data & Code Availability). Several non-seed hubs in this shared core are noteworthy both for their biological roles and because they are structurally interrogated later in the study. ACTN4, together with FLNB and FLNC, exemplifies the prominence of cytoskeletal crosslinkers that integrate adhesion, force transmission, and signalling. LMNB1, which pairs with lamin A/C, anchors chromatin to the nuclear periphery, reinforcing the nuclear-envelope theme that recurs across phenotypes. Similarly, the RPA1-3 complex exemplifies replication-stress adaptation, while EEF1D and the aminoacyl-tRNA synthetases QARS1 and EPRS1 link translation elongation and multi-synthetase complex function to proteostasis and stress regulation. TPR, a nuclear-pore basket scaffold, and UBE2I/UBC9, the central SUMO E2 conjugase, extend this nuclear quality-control axis by coupling transport and genome-stability checkpoints to SUMOylation. Proteostasis control is also underscored by VCP (p97/TERA), an AAA+ ATPase directing ER-associated degradation and organelle quality control. In addition, a recurrent signalling dimension is provided by the 14-3-3 adaptor family: YWHAB, YWHAG, and YWHAZ join their seeded counterparts in scaffolding phospho-proteins to regulate apoptosis, stress adaptation, and metabolic responses. Finally, VIM stands out as a type-III intermediate filament that interfaces with 14-3-3 proteins and organizes stress-adaptive cytoskeletal remodelling. Reactome analysis of the 127-gene set (Table 1) converged on stress-adaptation and proteostasis modules, including HSF1 activation and regulation of the heat-shock response, the calnexin/calreticulin ER-folding cycle, cytosolic tRNA aminoacylation, and initiation of nuclear-envelope reformation. We also observed signals across RAS-axis pathways (RAS-GTPase and RAS-GAP mutants; activation of RAS in immune cells), selenium-dependent amino-acid metabolism, and SARS-CoV-1/2 targeted host pathways, which often capture chaperone/ER-stress and MAPK components. At the GO level, the only biological-process term consistently shared was positive regulation of telomere maintenance via telomerase, aligning with the nuclear-envelope/genome-stability theme. Repeating the overlap analysis with downregulated PAOSA seeds yielded a highly consistent result: 107 genes were shared with the phenotype networks, of which 105 overlapped with the 127-gene set from the upregulated analysis. This near-identity indicates that both directions of expression change converge on the same molecular backbone. The few upregulated specific genes (e.g., IARS1, LARS1, QARS1, BAG6, GANAB, PDHA1, NDUFS3, PICALM, PRMT1, TXN, TOP1) reinforced themes of translation, proteostasis, and ER folding, whereas the two downregulated specific genes (ARF6, KRT31) mapped to peripheral immune and apoptosis pathways. Taken together, the under-expressed analysis functions mainly as a robustness check, while the 127-gene set serves as the principal core signature. The convergence of these results highlights that astrocytic oxidative stress responses map onto a conserved molecular architecture, simultaneously safeguarding proteostasis and genome integrity while embedding vulnerabilities in signalling and metabolic control that cut across neurodegenerative and cardiometabolic disease networks.

**Figure 2.**
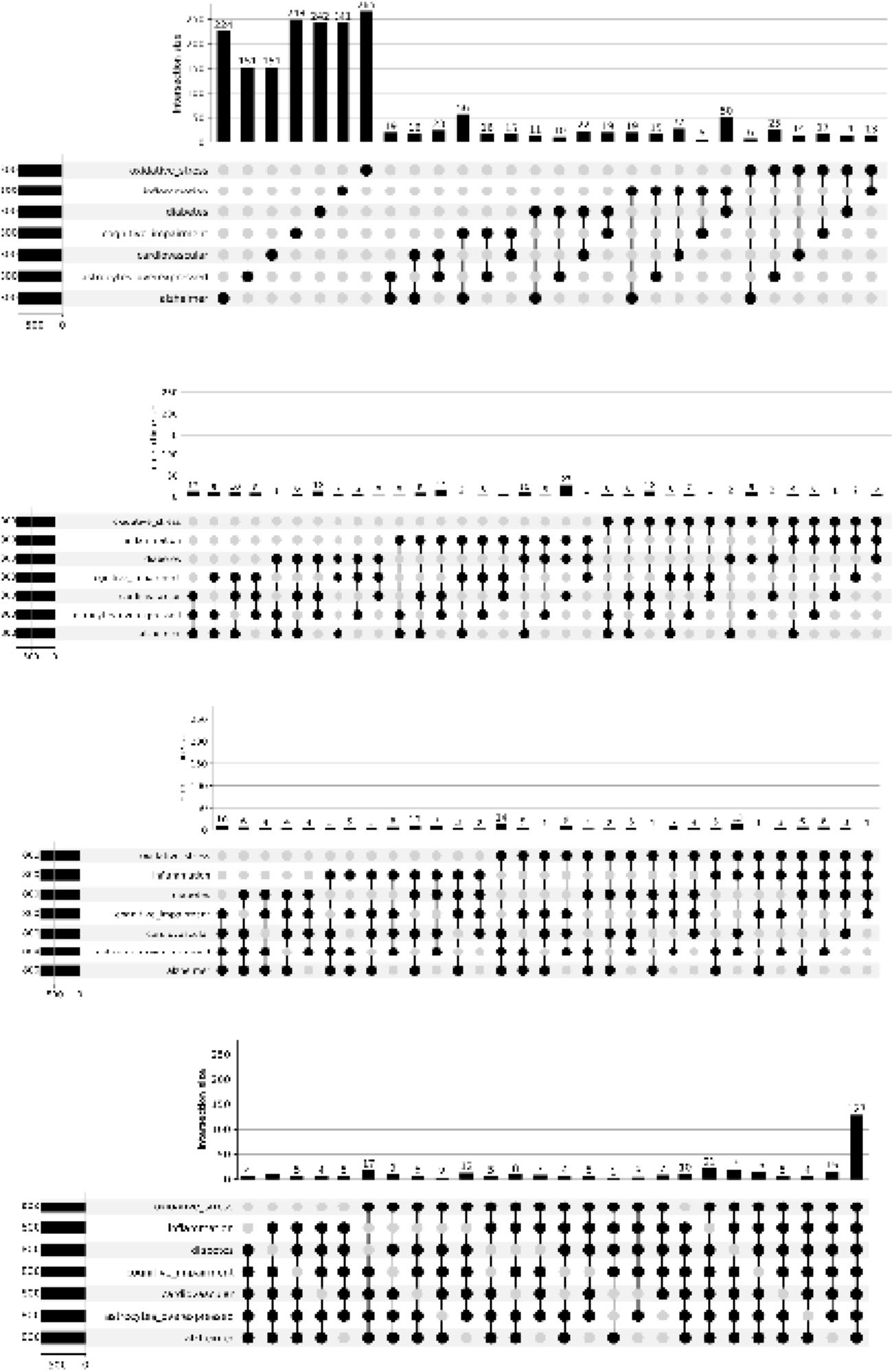
UpSet Plot of Shared Genes Across Phenotype Networks. The plot shows intersections among the top 5% of prioritized genes across seven networks: Alzheimer’s disease (AD), cardiovascular disease (CVD), type 2 diabetes (T2D), oxidative stress (OS), inflammation (I), cognitive impairment (CI), and astrocytes under oxidative stress (AO). Each row on the left represents one phenotype-specific gene list. Filled dots and connecting lines in the matrix indicate shared gene sets, while vertical bars above quantify the size of each intersection. The largest bar corresponds to the 127 genes present in all seven networks, defining a core molecular signature across aging-related conditions.

**Figure 3.**
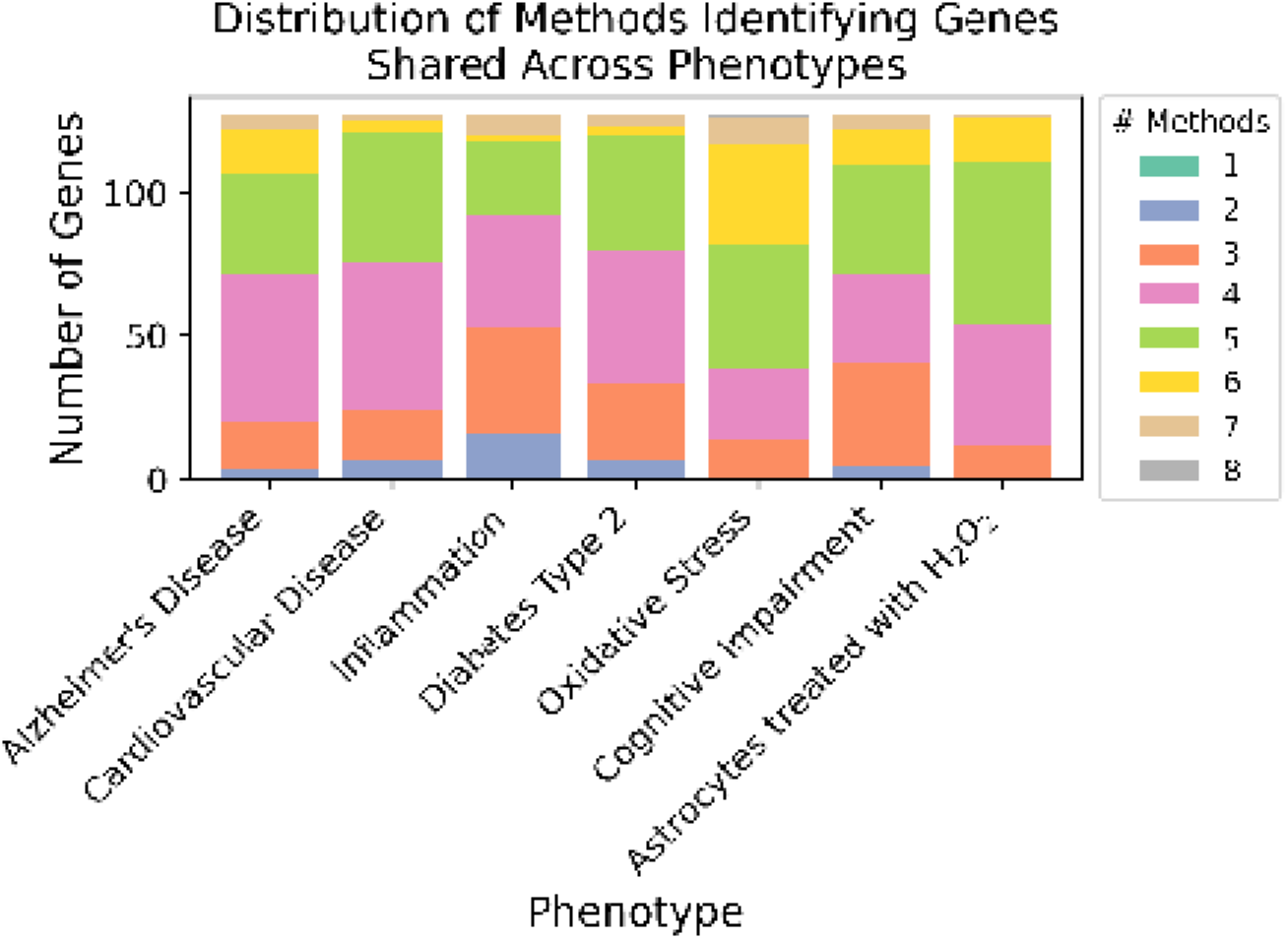
Stacked Bar Chart of Method Consensus per Phenotype. For each of the seven phenotypes (AD, CVD, T2D, OS, I, CI, AO), the stacked bar indicates how many of the eight topology-based ranking methods in GUILD identified each gene within the top 5% of prioritized candidates. The x-axis shows phenotypes, and the y-axis indicates the number of top-ranked genes. Colored segments represent the number of methods supporting each gene-disease predicted association (from 1 to 8). Higher segments reflect stronger consensus, with the top segments representing genes recognized by all eight methods. This visualization highlights the robustness of gene prioritization across analytic approaches.

**Table 1.** Reactome Pathway Enrichment Across Astrocyte and Disease Networks. This table lists Reactome pathways significantly overrepresented among the 127 genes shared by the astrocyte oxidative-stress response network (PAOSA) and the six disease-associated networks.

| Term name | Alzheimer's Disease | Cardiovascular Disease | Type 2 Diabetes | Oxidative stress | Inflammation | Cognitive Impairment | Astrocytes over | Astrocytes under |
| --- | --- | --- | --- | --- | --- | --- | --- | --- |
| Seenoamino acid metabolism | 2.84E-07 | 6.20E-08 | 2.21E-06 | 5.58E-09 | 2.84E-04 | 8.35E-04 | 2.41E-07 |  |
| Transcriptional and post-translational regulation of MITF-M expression and activity | 5.53E-05 | 6.53E-08 | 2.98E-08 | 1.88E-06 | 3.94E-05 | 9.59E-03 | 4.38E-03 | 2.11E-02 |
| Cytosolic tRNA aminoacylation | 1.22E-04 | 1.45E-06 | 2.65E-03 | 2.23E-08 | 2.55E-02 | 1.26E-02 | 3.94E-09 |  |
| HSF1 activation | 2.00E-04 | 7.40E-05 | 1.97E-03 | 7.60E-03 | 1.48E-03 | 4.37E-04 | 4.60E-05 | 1.43E-02 |
| SARS-CoV-1 targets host intracellular signalling and regulatory pathways | 4.85E-04 | 6.70E-08 | 4.01E-04 | 1.37E-04 | 2.38E-05 | 6.62E-05 | 1.56E-03 | 4.03E-03 |
| Signaling by phosphorylated juxtamembrane, extracellular and kinase domain K17 mutants | 5.40E-04 | 4.66E-03 | 4.11E-02 | 2.89E-02 | 3.08E-02 | 4.34E-03 | 1.19E-03 | 4.61E-02 |
| SARS-CoV-2 targets host intracellular signalling and regulatory pathways | 1.07E-03 | 3.80E-04 | 9.40E-05 | 4.03E-04 | 6.01E-03 | 1.95E-03 | 4.63E-03 | 1.02E-03 |
| Initiation of Nuclear Envelope (NE) Reformation | 2.38E-03 | 3.27E-05 | 1.13E-02 | 1.60E-05 | 7.81E-03 | 2.20E-05 | 1.19E-03 | 4.61E-02 |
| Signaling by RAS GAP mutants | 1.01E-02 | 4.55E-03 | 1.13E-02 | 6.56E-03 | 7.52E-03 | 1.26E-02 | 6.43E-03 | 1.23E-02 |
| Regulation of HSF1-mediated heat shock response | 1.65E-02 | 5.20E-05 | 2.76E-03 | 1.26E-06 | 1.27E-02 | 8.35E-04 | 4.22E-04 | 3.53E-03 |
| Activation of BAD and translocation to mitochondria | 1.71E-02 | 8.08E-03 | 4.01E-04 | 1.63E-03 | 2.47E-03 | 2.05E-02 |  | 2.11E-02 |
| CD209 (DC-SIGN) signaling | 1.71E-02 | 1.90E-03 | 4.70E-03 | 4.55E-02 | 3.60E-03 | 2.04E-02 |  | 2.11E-02 |
| EGFR Transactivation by Gasrin | 2.60E-02 | 1.45E-02 | 4.70E-03 | 1.55E-04 | 3.59E-03 | 6.61E-03 |  | 6.12E-03 |
| Calnexin/calreticulin cycle | 4.49E-02 | 2.12E-03 | 4.90E-02 | 2.97E-03 | 3.59E-02 | 4.10E-02 | 3.63E-02 |  |
| Activation of RAS in B cells | 4.49E-02 | 2.52E-02 | 4.90E-02 | 3.65E-02 | 3.59E-02 | 4.10E-02 | 3.63E-02 |  |

## 3. Structural analysis of interactions

We next explored the potential structural impacts of the set of common prioritized network interactions, especially in the context of known disease-associated mutations. Some of them have already been associated with some of the studied pathologies, but the majority had not been associated with them, and none was known to be associated with all. For the interaction edges connecting the top 5% of prioritized genes, we queried CM2D3 for structural models of those protein pairs and retrieved ∼200 curated complexes, built by template-based modelling when homologs were available and by data-driven docking for novel pairs. We then mapped ClinVar pathogenic/likely pathogenic missense variants onto predicted binding interfaces and used FoldX to estimate the change in binding free energy (ΔΔG = mutant − wild-type). By convention, positive ΔΔG indicates a destabilizing mutation (weaker binding), whereas negative ΔΔG indicates a stabilizing mutation (stronger or hyperstable binding). This variant centric, structure informed approach complements network level prioritization and mirrors emerging frameworks that map disease variants to modelled interactomes to generate mechanistic hypotheses (Figure 4).

**Figure 4.**
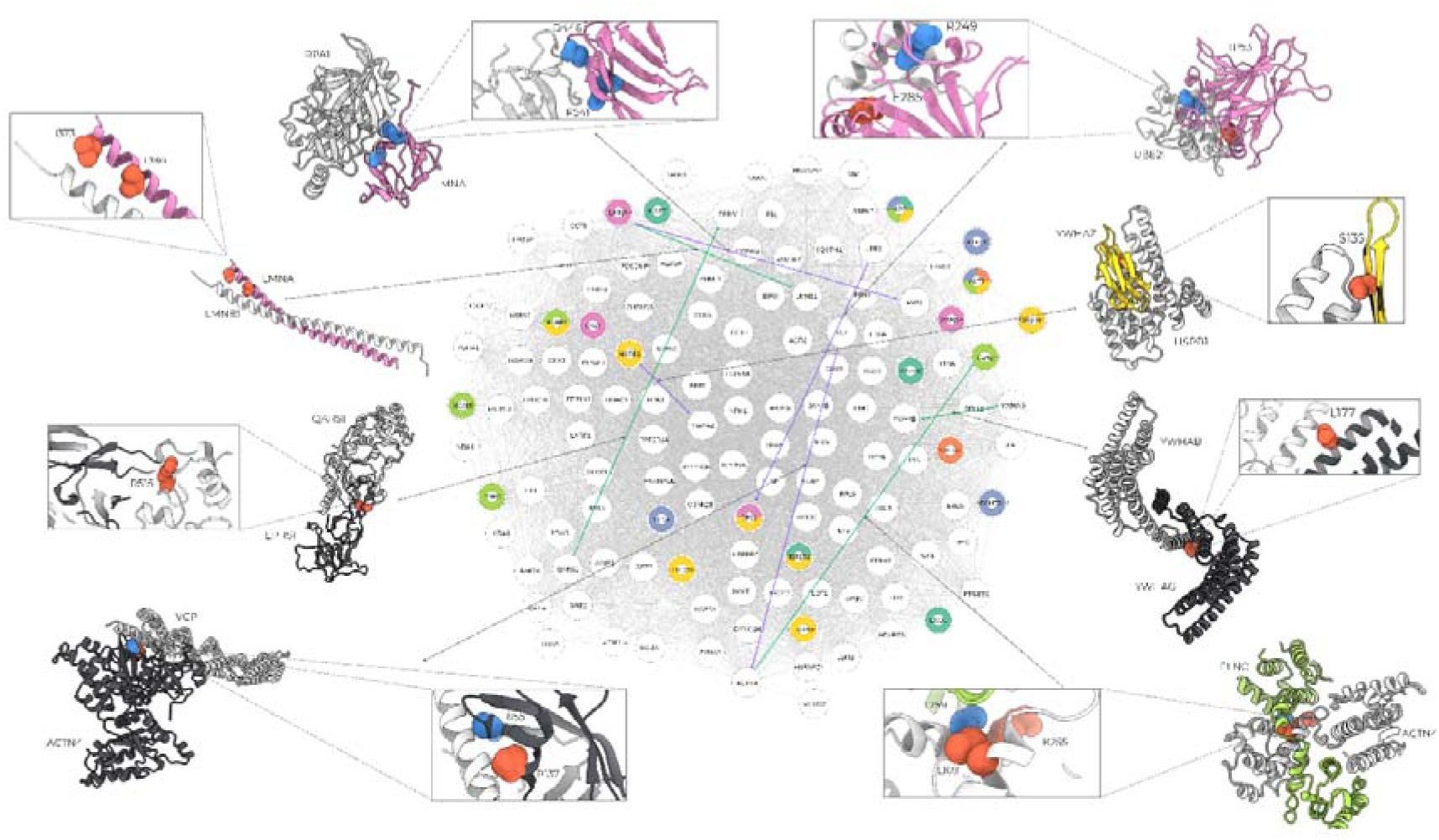
Cross-phenotype 127-gene interactome with CM2D3 structural support and variant effects. Cytoscape network of the 127 genes shared across astrocyte oxidative stress and six aging-related pathophenotypes. Node colors mark seed membership to each pathophenotype (see legend below); white nodes are non-seed genes within the shared core. Multi-phenotype seeds are rendered as pie-chart nodes with segment colors indicating phenotype memberships. Edges denote PPIs with structural models from CM2D3: green = homology/template-based complexes; violet = docking-derived complexes. Structural insets (rendered in Chimera) show representative interfaces; zoomed residues indicate missense positions whose mutations alter binding energies computed with FoldX (ΔΔG_interface = mutant − wild-type). Residues depicted with atom-spheres in red indicate destabilizing mutations and hyperstabilizing mutations are shown in blue. Phenotype color legend: Pink = Diabetes, Orange = Cardiovascular, Green = Inflammation, Turquoise = Alzheimer, Yellow = Oxidative Stress, Blue = Cognitive Impairment

Across CM2D3’s template-based models, many disease variants were predicted to weaken protein–protein interactions (PPIs), consistent with loss-of-interaction mechanisms (Table 2).

**Table 2.**
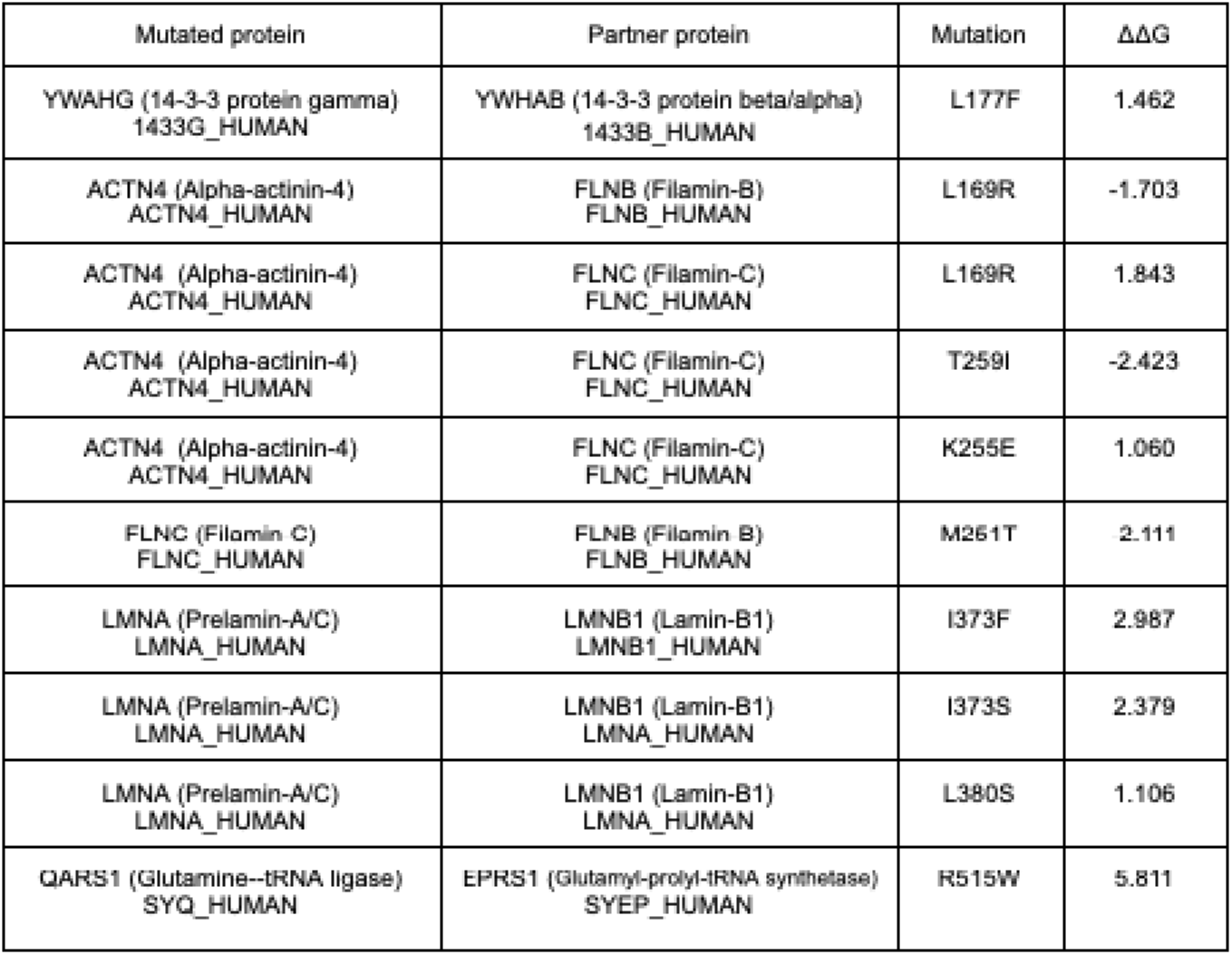
Structural perturbations at PPI interfaces in comparative (homology-based) models. Summary of key pathogenic missense variants mapped onto interfaces of 3D complexes obtained via CM2D3 template-based modelling. For each mutated protein and its binding partner, the specific amino-acid substitution is listed along with the FoldX-calculated change in binding free energy (ΔΔG = mutant – wild-type) in kcal/mol. Positive ΔΔG values indicate destabilizing mutations (loss of interaction affinity), whereas negative values indicate stabilizing (gain-of-binding) effects.

A representative example is the nuclear lamin heterodimer (LMNA–LMNB1), where multiple lamin-A missense variants associated with laminopathies cluster at the interface. LMNA I373F and I373S showed among the largest ΔΔG increases (> +2.3 kcal/mol), indicating substantially reduced heterodimer affinity; additional interface variants (R377H, L380S) also yielded positive ΔΔG (+0.5–1.1 kcal/mol). Not all substitutions were destabilizing: LMNA E358K slightly decreased ΔΔG (≈ −0.4 kcal/mol), suggesting a modest *gain* of binding that could stiffen the lamina and impair its dynamics. Together, these results support a model in which both loss- and gain-of-binding at A–B lamin contacts can perturb nuclear mechanics, chromatin tethering, and stress responses linked to aging phenotypes^102,103^. A second prominent module involves the actin cytoskeleton, particularly α-actinin-4 (ACTN4) complexes with filamins (FLNB/FLNC). Variants originally linked to kidney disease or myopathy often fell directly at the ACTN4–filamin interface^104,105^. For example, ACTN4 K255E, a familial FSGS mutation^106^, was predicted to weaken binding (ΔΔG ≈ +0.8 to +1.1 kcal/mol across ACTN4–FLNB/FLNC), potentially compromising actin crosslinking and mechanotransduction^107^. Notably, the impact of ACTN4 L169R depended on partner: it strengthened ACTN4–FLNB (ΔΔG ≈ −1.7 kcal/mol) but destabilized ACTN4–FLNC (ΔΔG ≈ +1.8 kcal/mol), illustrating context-specific effects. Conversely, ACTN4 T259I strongly stabilized ACTN4–FLNC (ΔΔG ≈ −2.4 kcal/mol), suggesting pathologically tight binding.

We also observed interface-sensitive effects on 14-3-3 dimerization^108^. In models of YWHAG/YWHAZ dimers, a L177F/L178F-like substitution yielded a moderate ΔΔG increase (≈ +1.4 kcal/mol), consistent with weakened dimer formation^109,110^ and potentially reduced availability of 14-3-3 scaffolds for phospho-client signalling complexes; by contrast, YWHAZ V180A produced only a mild effect (+0.3–0.6 kcal/mol). Given the centrality of 14-3-3 in stress signalling, even modest dimer weakening could reverberate across apoptotic and metabolic pathways^111^.

Finally, within the multi-aminoacyl tRNA synthetase complex (MSC), a pathogenic QARS1 R515W variant mapped directly to the QARS1–EPRS1 interface and produced one of the largest ΔΔG values we observed (≈ +5.8 kcal/mol), implying near-complete disruption^112^. Dissociation of MSC components, specifically EPRS1, drives the interferon-γ/TLR-responsive GAIT pathway, which represses translation of inflammation-related mRNAs, linking a single variant to proteostasis pressure and chronic inflammation^113,114^.

Further analyses on docking-based complexes revealed diverse, partner-specific effects (Table 3). In LMNA–RPA1, LMNA D446N and R541G markedly strengthened binding (ΔΔG ≈ −18.7 and −10.2 kcal/mol), predicting aberrantly stable assemblies with potential consequences for nuclear stability and aging phenotypes. By comparison, LMNA variants in complexes with EEF1D and TPR had more moderate effects. For TP53–UBE2I (UBC9), we observed both directions of change: TP53 R249S significantly strengthened binding (ΔΔG ≈ −3.4 kcal/mol), potentially altering SUMOylation dynamics, whereas TP53 E285K destabilized the complex (ΔΔG ≈ +2.3 kcal/mol). The VCP (p97; formerly TERA)–ACTN4 interface was especially sensitive: VCP R155G dramatically stabilized binding (ΔΔG ≈ −23.8 kcal/mol), while nearby P137L strongly destabilized it (ΔΔG ≈ +5.5 kcal/mol). Stress-response chaperone interfaces were similarly variant-sensitive: HSPB1 S135Y severely weakened HSPB1–14-3-3ζ (ΔΔG ≈ +9.6 kcal/mol), whereas R140G modestly strengthened HSPB1–14-3-3β (ΔΔG ≈ −3.4 kcal/mol).

**Table 3.**
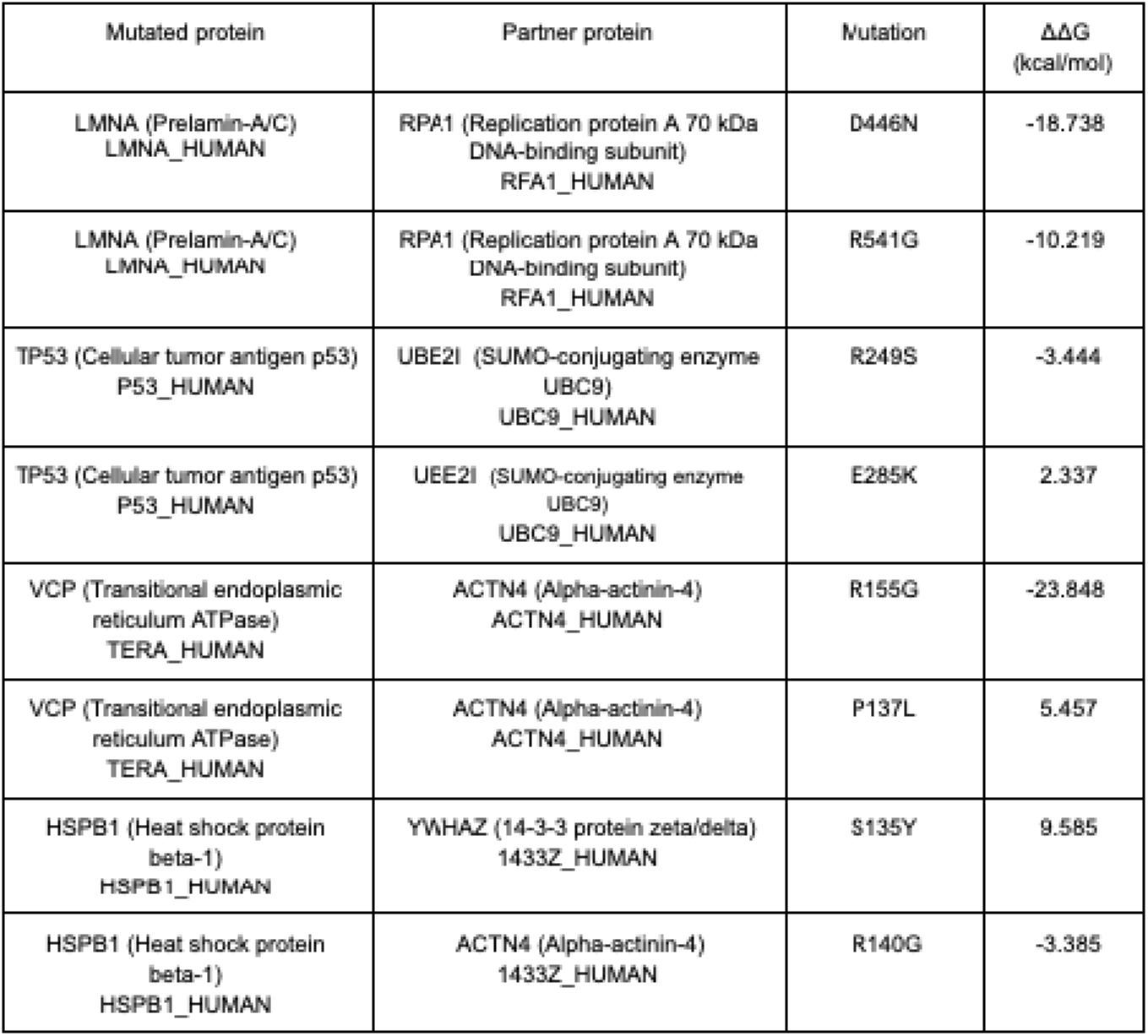
Structural perturbations at PPI interfaces in docking-derived models. Summary of key pathogenic missense variants mapped onto interfaces of complexes generated by ab initio docking. Each row gives the mutated protein and its docking partner, the residue change, and the FoldX ΔΔG (kcal/mol). As before, positive ΔΔG denotes predicted weakening of the interaction, and negative ΔΔG denotes predicted strengthening.

In parallel, we leveraged PRISM to obtain interface predictions for selected high-priority interactions among the top-ranked overlap genes, focusing on pairs not structurally covered by CM2D3. PRISM identifies candidate complexes by matching surface patches of the target proteins to complementary sides of experimentally resolved template interfaces and ranks them by a global interaction-energy score, with more negative values indicating more favourable interfaces. We interpreted these PRISM-derived complexes alongside the CM2D3/FoldX ΔΔ*G_interface* results as orthogonal structural support for specific subnetworks. Within this subset, two motifs were particularly prominent: (i) a 14-3-3 signalling hub, in which YWHAZ engages TP53, AKT1, VIM, and STK11-containing assemblies, and (ii) an APP–proteostasis axis connecting APP and γ-secretase conformers to VCP/p97. The full set of PRISM predictions, together with the CM2D3 models and ClinVar-mapped mutations, is available in the GitHub repository (see Data & Code Availability).

## Discussion

This work links a human astrocyte oxidative-stress model to disease mechanisms by integrating transcriptomics, network-based prioritization, and structure-guided variant interpretation. Three main conclusions emerge. First, sublethal oxidative stress rewires astrocytes toward a state that simultaneously boosts proteostasis safeguards and translational control while inhibiting chromatin/transcriptional programs, an adaptive but potentially brittle configuration if stress persists. Second, when embedded in the human interactome and projected onto aging-related phenotypes (AD, CVD, T2D, CI, inflammation, OS), these astrocytic signals converge on a shared 127-gene core enriched for proteostasis, DNA repair, mitochondrial regulation, and telomere/nuclear-envelope maintenance. Third, structural modelling, combining CM2D3 complexes with PRISM-derived complexes for high-priority pairs lacking templates, pinpoints variants likely to perturb these systems and highlights structurally plausible interfaces, offering mechanistic routes from network hubs to cellular failure modes.

The phenotype-stratified subnetworks capture canonical disease axes but also reveal their overlap within astrocytes. AD- and CI-focused analyses highlight ER-associated folding/clearance, insulin/IGF signalling, synaptic support, and nitric-oxide/cerebrovascular coupling features that are also represented in the CVD network via endothelial function, lipid handling, and circadian regulation. The T2D subnetwork reinforces immunometabolic crosstalk (INSR/IRS/AKT pathways with NF-κB and oxidative defences), while the inflammation and OS networks embed these same regulators in stress-adaptation circuits. That all six disease subnetworks intersect with the astrocyte oxidative-stress network in an unexpectedly large 127-gene core argues for a conserved vulnerability architecture. Notably, recurring non-seed hubs, TGOLN2 (secretory sorting), the RPA1–3 complex (replication/repair), and LMNA (nuclear lamina), bridge modules that are often treated separately (secretory traffic, genome maintenance, and mechanotransduction), offering a parsimonious explanation for the observed comorbidity between neurodegenerative and cardiovascular conditions in aging. A salient theme is envelope/genome stability: enrichment for positive regulation of telomere maintenance and initiation of nuclear-envelope reformation dovetails with the prominence of LMNA and replication-stress nodes (RPA, ATR/ERCC factors). In parallel, proteostasis emerges as a central pillar (HSF1 activation, calnexin/calreticulin cycle, cytosolic tRNA aminoacylation), consistent with the transcriptional up-shift of chaperone and translational quality-control genes. Together, these results support a model in which aging astrocytes attempt to preserve protein and genome integrity under chronic redox pressure, but in doing so reprogram signalling and trafficking in ways that can propagate dysfunction to the neurovascular unit and systemic metabolism. Mapping ClinVar missense variants onto CM2D3 complexes showed that many disease-linked substitutions likely weaken PPIs (positive ΔΔG), but “gain of binding” also appears pathogenic in specific contexts. The LMNA-LMNB1 heterodimer illustrates both behaviors: several lamin-A variants markedly destabilize the interface, while others modestly stabilize it, implying either lamina fragility or pathological stiffening could impair mechanotransduction, DNA-damage responses, and stress resilience. In the actin–adhesion module, ACTN4 substitutions exert partner-specific effects on filamin binding, predicting altered crosslinking and force transmission processes relevant to astrocyte morphology, perivascular endfoot integrity, and BBB support. For 14-3-3 scaffolds, dimer-weakening substitutions may reduce the availability of client-binding platforms that coordinate phospho-signalling under stress, with ramifications for apoptosis and metabolic control. Within the multi-aminoacyl-tRNA synthetase complex, the large ΔΔG predicted for QARS1-EPRS1 disruption suggests a path to proteostasis collapse and aberrant inflammatory translation (GAIT), providing a direct link between redox stress and chronic inflammation.

PRISM analyses supplied orthogonal support in the subset of high-priority interactions lacking CM2D3 complexes. Several hubs repeatedly achieved favorable interface matches with strongly negative global energies, including: (i) a 14-3-3 (YWHAZ/YWHAG) hub engaging TP53, AKT1, VIM, and STK11-containing assemblies; (ii) an APP–proteostasis axis connecting APP/γ-secretase conformers to VCP/p97. Mechanistically, these complexes tie oxidative stress and proteostasis (APP–VCP), checkpoint and survival signalling (p53–14-3-3ζ, AKT1–14-3-3) to the shared astrocytic vulnerability backbone. The cross-phenotype core suggests that biomarkers and interventions targeting common nodes may yield benefits across neurodegenerative and cardiometabolic spectra. Candidate biomarkers include secretory-pathway components (e.g., TGOLN2-dependent cargo), lamina-derived fragments or post-translational signatures, and DNA-damage/repair markers (RPA-associated). Therapeutically, three leverage points emerge: (1) proteostasis reinforcement (HSF1/UPR tuning, selective autophagy/ERAD enhancement); (2) nuclear-envelope/genome-stability support (modulating lamin dynamics, DDR signalling, oxidative DNA-repair capacity); and (3) scaffolded signalling control (stabilizing 14-3-3 dimer/client interactions or correcting hyper-stabilized/unstable variant interfaces). The APP–VCP connection specifically suggests testing p97 modulators or trafficking-chaperone strategies for effects on APP/γ-secretase routing in astrocytes, with dual readouts in amyloid processing and vascular/endothelial crosstalk.

## Data & Code Availability

Data and code are publicly available at https://github.com/abotlp/Astrocytes_paper. Within the repository, all analysis outputs are organized under *Astrocytes_project/GUILDifyTools*-*main/astrocytes*. For each phenotype (alzheimer, cardiovascular, cognitive_impairment, diabetes, inflammation, oxidative_stress) and for the astrocyte runs (*astrocytes_overexpressed, astrocytes_underexpressed*), the seed genes are listed in seeds.txt, the consensus GUILD ranking is provided in guild_scores.txt, and functional enrichment tables (GO BP/MF and *Reactom*e) are in *enrichment.GObp.*, enrichment.GOmf.*, and enrichment.Reactome.\** (where * is any supplementary suffix). Crosslzlnetwork overlap results are in *network_overlap_over5* and *network_overlap_under5*, including the shared gene lists (*protein_overlap_results.txt*), pathway overlaps (*function_overlap_results.txt*), and the overlap subnetworks (*subnetwork_genes.sif*). All analyses are built on the BIANAlzlderived human protein-protein interaction network available in *Astrocytes_project/BIANA_phy*. The complete set of mutation analyses referenced in the paper (*FoldX* ΔΔ*G* summaries and ClinVar mappings) is available under *Astrocytes_project/ddg_analysis*.

## Acknowledgements

This work was funded by the Spanish Institute of Health Carlos III by project reference AC20/00009-FEDER/UE and ERANET ERA-CVD_JTC2020-015. This work was also supported by the Spanish Ministry of Science and Innovation and Agencia Estatal de Investigación plus FEDER Funds through grants PID2023-150068OB-I00 funded by MICIU/AEI/10.13039/501100011033 and by ‘ERDF A way of making Europe’ (BO) and PID2023-149767OB-I00 funded by MICIU/AEI/10.13039/501100011033 and by ’ERDF A way of making Europe’ (FJM), and, ‘Unidad de Excelencia María de Maeztu’ CEX2024-001431-M, funded by MICIU/AEI/10.13039/501100011033. This project was funded in part by TUBITAK Research GrantNo: 220N252 (AG). S. A. acknowledges the support provided by Erasmus+ for the study program. The authors thank the Scientific Computing Core Facility (MELIS-UPF).

---

**Supplementary Table 1.**
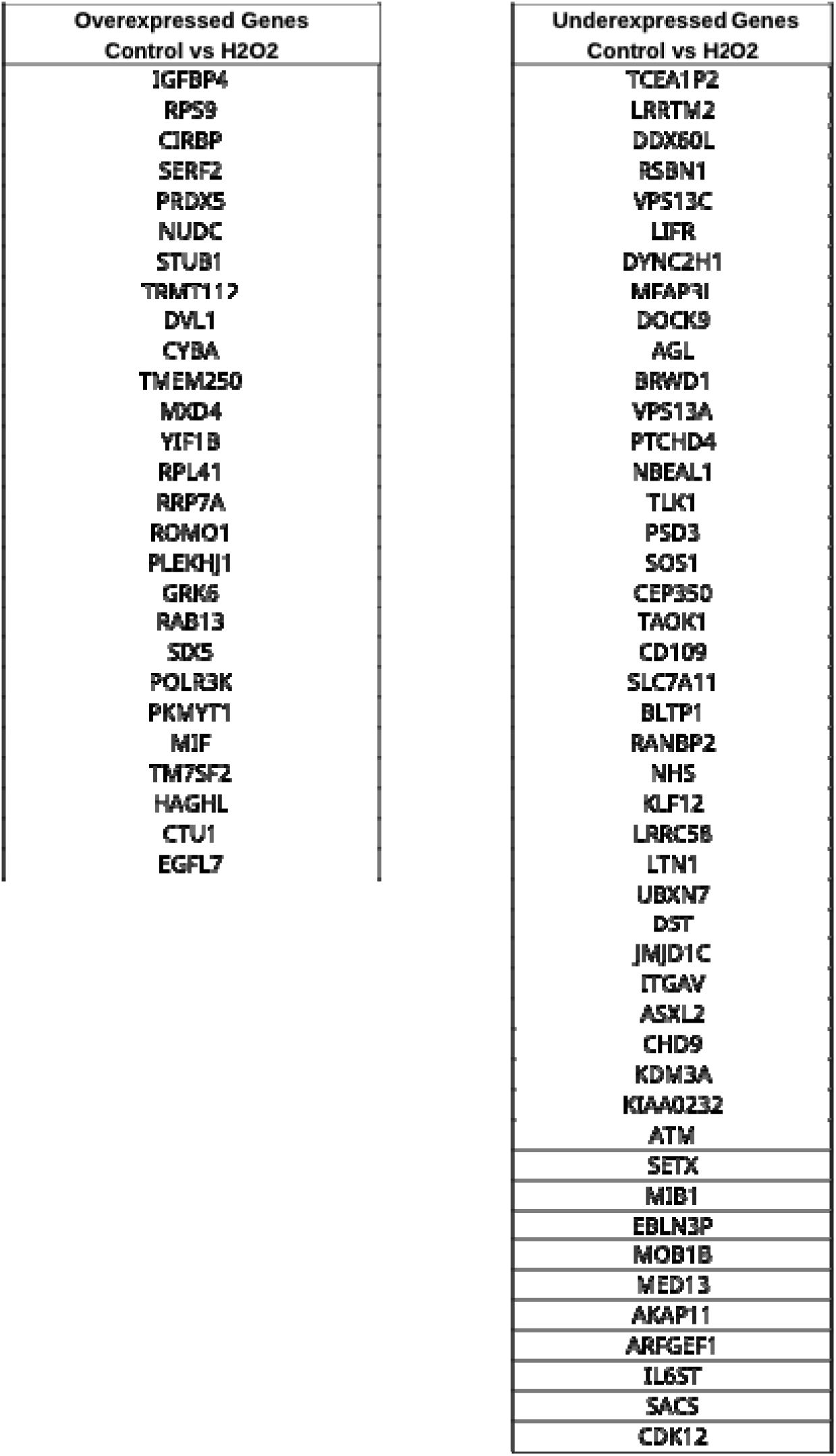
Downregulated and upregulated genes in astrocytes after stress by H_2_O_2_.

## Notes

https://github.com/abotlp/Astrocytes_paper

### Competing Interest Statement

The authors have declared no competing interest.

